# ACE-2-like Enzymatic Activity in Anti-SARS-CoV-2 Spike Protein Monoclonal Antibodies

**DOI:** 10.64898/2026.02.02.703244

**Authors:** Yufeng Song, Frances Mehl, Tom No, Lauren Livingston, Juan Sebastian Quintero Barbosa, Jun Hayashi, Ginette Serrero, Pamela Schoppee Bortz, Jeffrey M Wilson, James E Crowe, David D Ho, Michael T Yin, Joshua Tan, Steven L Zeichner

## Abstract

Many people with acute COVID-19 have clinical disease not clearly attributable to viral replication and many COVID-19 convalescents are affected by post-acute sequelae of COVID-19 (PASC, or long COVID, LC). LC has severely affected public health and economies worldwide. Features of LC including blood pressure dysregulation, coagulopathies, high levels of inflammation, and neuropsychiatric complaints. The mechanisms responsible for the pathogenesis of some of COVID-19’s clinical features and LC have not been well established. The host cell receptor for SARS-CoV-2 is human angiotensin converting enzyme 2 (ACE2), which binds the SARS-CoV-2 spike protein receptor-binding domain (RBD) to initiate infection. We hypothesized that some people may produce anti-RBD antibodies that sufficiently resemble ACE2 structure to have ACE2-like catalytic activity after infection. Those antibodies, ACE2-like abzymes, may contribute to the pathogenesis of LC. Our previous studies showed that ACE2-like activity was associated with immunoglobulin in some acute and convalescent COVID-19 patients. ACE2-like catalytic activity correlated with blood pressure changes following a moderate exercise challenge in people convalescing from COVID-19. To further establish that ACE2-like activity could be attributed to antibodies, we screened human monoclonal antibodies (mAbs) against SARS-CoV-2 spike protein from 3 different research centers and others purchased from a commercial source for ACE2-like catalytic activity. We identified 4 human monoclonal antibodies with ACE2-like catalytic activity. The ACE2-like catalytic activity of these mAbs was not inhibited by MLN-4760, a compound that inhibits native human ACE2 catalytic activity, nor by EDTA, unlike native ACE2, a Zinc metalloprotease, but was inhibited by an overlapping pool of spike peptides. Enzyme kinetic studies showed that the mAbs had substantially lower Vmax and *Km* values than native ACE2. The data therefore suggested that the antibodies cleave ACE2 substrate via a different mechanism than native ACE2. The identification of specific mAbs with ACE2-like catalytic activity supports the hypothesis that antibodies induced by SARS-CoV-2 infection could help mediate the pathogenesis of COVID-19 and LC, and more generally, the hypothesis that catalytic antibodies induced by infectious agents can contribute to disease pathogenesis.

## Introduction

COVID-19 and Post-Acute Sequelae of COVID-19 (PASC, or Long COVID, LC), as reviewed in ^1–3^, can affect up to 7% of COVID-19 patients^4^. LC patients can have many unusual symptoms such as dizziness, fatigue, malaise, “brain fog”, alterations in smell or taste, abnormal movements, libido changes, and other neuropsychiatric symptoms, GI symptoms, cardiovascular complaints such as palpitations, blood pressure dysregulation, inflammation, coagulopathies, cough, and chest pain^5^ ^6, 7^. While the clinical features, laboratory findings, and epidemiology of LC have been extensively described, clear primary causal etiologies have yet to be determined ^8^.

Antibodies with catalytic activity, also called “abzymes” have been known for some time ^9, 10^. Investigators initially produced abzymes by generating anti-idiotypic antibodies against antibodies targeting enzymes’ active sites^11, 12^. In some instances, the anti-idiotypic antibodies had sufficient resemblance to the original enzyme’s active site so as to exhibit catalytic activity similar to that of the initially targeted enzyme^13, 14^. However, further research showed that the abzymes had decreased substrate specificity and less favorable enzyme constants compared to the targeted enzymes (*e.g., k*cat/*Km* values ∼10^2^ - 10^4^ s^-1^•M s^-1^ ^15^ vs. *k*cat/*Km* values of ∼10^5^ s^-1^•M s^-1^)^16^ ^12, 13, 17^. These limitations led to a decline in enthusiasm for the use of abzymes in biotechnological applications.

Antibodies are traditionally understood as high affinity binding molecules. However, a subset of antibodies with intrinsic catalytic activity, commonly referred to as natural abzymes, has been described in a variety of immune contexts, including autoimmune disease, chronic viral infection, and exposure to persistent antigens^18–20^. Such antibodies typically exhibit low catalytic turnover relative to canonical enzymes, have heterogeneous substrate preferences, and occur at low frequencies within polyclonal repertoires^18, 20–22^. While the catalytic efficiency of such abzymes is lower than typical enzymes, their biological significance lies in their persistence, the potentially high concentration of immunoglobulin molecules in the circulation, and systemic distribution^23^. With respect to SARS-CoV-2, anti-spike protein antibodies have been observed in the nanomolar range ^24^.

Abzymes have been shown to exert measurable biochemical activity and, in certain contexts, have been implicated in immune-mediated modulation of host physiology^18, 20, 25, 26^. Given that homeostatic regulatory mechanisms are finely balanced, low-level hydrolysis of regulatory peptide, like angiotensin II, could shift the homeostatic “proteolytic tone“^27^, although the short half-life of regulatory peptides, like angiotensin II (less than 1 minute) ^28^ make meaningful *in vivo* measurements difficult.

In an attempt to identify potential mechanisms responsible for the unusual features of COVID-19 and LC, we considered that many of these aberrant clinical features could be viewed as consequences of pathologic derangements of proteolytic regulatory cascades, for example, blood pressure regulation (the renin-angiotensin system), inflammation (C’, kallikrein-kinin)^29–31^, and coagulation. For LC, some most recent studies found the kinetics of symptom development would be consistent with antibody production, rather than the pathology directly produced by viral replication^32, 33^. In addition, a recent report showed that immunoglobulins transferred from some LC patients to mice can induce symptoms, further suggesting that immunoglobulin may help mediate some LC symptoms^34^.

While the benefits of COVID-19 vaccination vastly outweigh the risks, the fact that a very small, but still non-zero, fraction of patients who received SARS-CoV-2 vaccines had symptoms including hyperinflammation, blood pressure dysregulation, or coagulopathies ^35–37^ further suggests that, in some cases, an antibody response could play a role in pathogenesis of LC, as opposed to lasting sequelae of active viral replication.

Angiotensin Converting Enzyme 2 (ACE2) on the host cell surface serves as the principal receptor for the SARS-CoV-2 Receptor Binding Domain (RBD). Angiotensin Converting Enzyme (ACE) catalyzes the cleavage of angiotensin I to angiotensin II, which acts as vasoconstrictor to increase blood pressure ^38^. ACE2 cleaves angiotensin II to produce angiotensin 1-7, which has vasodilatory activity that lowers blood pressure, an activity that ACE2 counteracts the activity of ACE in the renin–angiotensin system by converting angiotensinlJII to angiotensinlZl(1–7), opposing ACElZldriven signaling^39^. While interactions between the SARS-CoV-2 Spike protein and the human ACE2 receptor are well-documented at the binding interface, the functional consequences of this interaction remain under-explored^40^. We hypothesized that the pathogenesis of some of the unusual features of LC might be caused by the induction of antibodies against the SARS-CoV-2 Spike protein (S) that resembled the structure of ACE2 closely enough to have ACE2-like catalytic activity.

In our previous studies^41, 42^, we found that some patients’ plasma with ACE2-like activity did not require divalent cations, unlike native ACE2, a Zinc metalloprotease. This activity co-depleted with immunoglobulin and was inhibited by a tiled overlapping RBD peptide pool. These findings suggested that the ACE2-like activity observed in patients’ plasma differed from native ACE2 and was the result of anti-RBD immunoglobulins exhibiting ACE2-like activity.

Additionally, ACE-2 like catalytic activity found in plasma collected from convalescent patients correlated with changes in blood pressure following a moderate exercise challenge^41^, suggesting that the catalytic antibodies could have physiologic consequences.

However, to definitively establish that COVID-19 patients produce antibodies with catalytic activity, it is important to identify a pure monoclonal antibody derived from COVID-19 patients that exhibits the hypothesized catalytic activity. Here, we survey several sets of monoclonal antibodies (mAbs) from 4 different research centers.

We utilized a widely used surrogate substrate which enables sensitive, reallZltime detection of catalytic activity and is commonly employed in ACE2 activity and inhibitor screening assays ^43–45^ to monitor ACE2-like activity. While we found that most mAbs lacked catalytic activity, two mAbs from one set showed catalytic activity higher than the level from the standard positive control, and two others showed borderline detectable activity. These observations show that some patients develop antibodies with catalytic activity following SARS-CoV-2 infection, with potential clinical consequences, providing a biochemical basis for how a virus-induced immune response might transition into a chronic, autoimmune catalytic process.

## Materials and Methods

### Monoclonal Antibody Collections

We received 129 purified human anti-SARS-CoV-2 RBD mAbs, 1 purified anti-SARS-CoV-2 fusion peptide (FP) mAb, 7 human anti-SARS-CoV-2 RBD mAb hybridoma supernatants, 3 irrelevant mAb hybridoma supernatants, and 1 empty hybridoma culture media as a negative control from 4 different collaborator groups (Table 1). Some of the anti-RBD antibodies were further characterized for their ability to block or enhance ACE2 binding to SARS-CoV-2 spike protein. The hybridoma culture medium is 1/3 Medium E (Stem Cell Technologies) and defined 2/3 Dulbecco’s Modified Eagle Medium (DMEM) + 10 % fetal bovine serum (FBS).

**Table 1:** Summary of Monoclonal Antibody Characteristics Used in the Study.

| Source | Total<br>Number | Form | Characteristics | Number<br>with<br>Abzyme<br>Activity | Reference |
| --- | --- | --- | --- | --- | --- |
| Dr. James<br>Crowe,<br>Vanderbilt<br>University<br>Medical<br>Center | 2 | Purified | Human Anti<br>SARS-CoV-2<br>RBM | 0 | 46, 47 |
| Dr. Jun<br>Hayashi,<br>Precision<br>Antibody | 3 | Supernatant | Human Anti<br>SARS-CoV-2<br>Spike and RBD,<br>block RBD<br>binding to ACE2 | 2 |  |
|  | 4 | Supernatant | Human Anti<br>SARS-CoV-2<br>Spike and RBD,<br>enhance RBD<br>binding to ACE2 | 2 |  |
|  | 3 | Supernatant | Negative Controls | 0 |  |
| Dr. David Ho,<br>Columbia<br>University | 39 | Purified | Human Anti<br>SARS-CoV-2<br>RBD | 0 | 49, 59, 97 |
| Dr. Joshua Tan at NIH | 88 | Purified | Human Anti SARS-CoV-2 RBD | 0 | <sup>52</sup> |
|  | 1 | Purified | Human Anti SARS-CoV-2 FP (Negative Control) | 0 | <sup>51</sup> |

### Monoclonal Antibodies for Screening

All antibodies were assessed for binding to SARS-CoV-2 RBD (with the exceptions of the negative control mAbs) as described^46–52^. The mAbs purchased from Precision Antibodies targeting the RBD (but not those from other sources) were further screened for interactions with ACE2. This screening used (spike) S and (receptor binding domain) RBD recombinant proteins with HIS tag and an ACE2 human Fc fusion protein. Ni-NTA plates were used to capture both S and RBD proteins. ACE2 binding to both S and RBD proteins was confirmed to ensure that both S and RBD retained the ability to bind ACE2. The S protein captured on Ni-NTA plates was used for the initial screening. All the clones that showed binding to S-protein were subsequently screened for specific RBD binding using RBD-captured Ni-NTA plates. ACE2 was added to the plate to see if any of the anti-RBD antibodies blocked ACE2 binding to the S or RBD proteins captured on the plate. The binding of ACE2 was detected using HRP-anti-human Fc.

### Angiotensin II Converting Enzyme (ACE2)-like Activity Assays

For the ACE2 activity assay, 25 µg purified antibody or 50 µL hybridoma supernatant before and after immunoglobulin depletion was combined with 50 µM MCA-APK(Dnp) fluorogenic ACE2 substrate (AAT Bioquest, Pleasanton CA. Cat. #1355) in assay buffer, 50 mM MES assay buffer (Thermo Fisher, Waltham MA. Cat. # J62231.AK) supplemented with 300 mM NaCl and 10 µM ZnCl_2_, in a non-binding black bottom plate (Corning, Glendale AZ. Cat. #3991) ^53, 54^ with total 100µL reaction volume^55^. Empty hybridoma culture media was used as a negative control. Antibody concentration in hybridoma supernatant was determined by IgG ELISA (Table 4) and antibody concentration in substrate catalytic reaction is shown in Table 3. Fluorescence levels were measured with excitation/emission wavelengths 320/405nm and a 325 nm emission filter on the SpectraMax® M5 multi-mode microplate reader (Molecular Devices, San Jose CA.)^54^. Measurements were taken every five minutes for 16 hours, while the plate was kept at a constant temperature of 37°C. 5 mM EDTA was added to the hybridoma culture supernatants to inhibit the ACE2 present in 10% FBS medium^56^. Specificity of ACE2 substrate cleavage was determined by parallel comparisons of samples with 50 µM OMNIMMP fluorogenic control substrate (Enzo Life Sciences, Farmingdale, NY. Cat. #MML-P127), MCA-Pro-Leu-OH, according to previous studies^41, 42, 57^.

### ACE2 Substrate Peptide Cleavage Competition Inhibition by SARS-CoV-2 Spike RBD Peptide Pools

Twelve peptides (SB-Peptide, SmartBioscience SAS, France) covering the SARS-CoV-2 RBD (including RBM) were reconstituted in ddH2O or DMSO to a concentration of 10 µg/µL (Table 2) ^41, 42^. The peptides were combined in equal amounts. The ACE2 activity assay was set up as described above, and 2.4 µL of the peptide pool was added so that there was 2 µg of each peptide per well. 2.4 µL of peptide diluent (50% ddH_2_O and 50% DMSO) was added to other wells as a control.

**Table 2:**
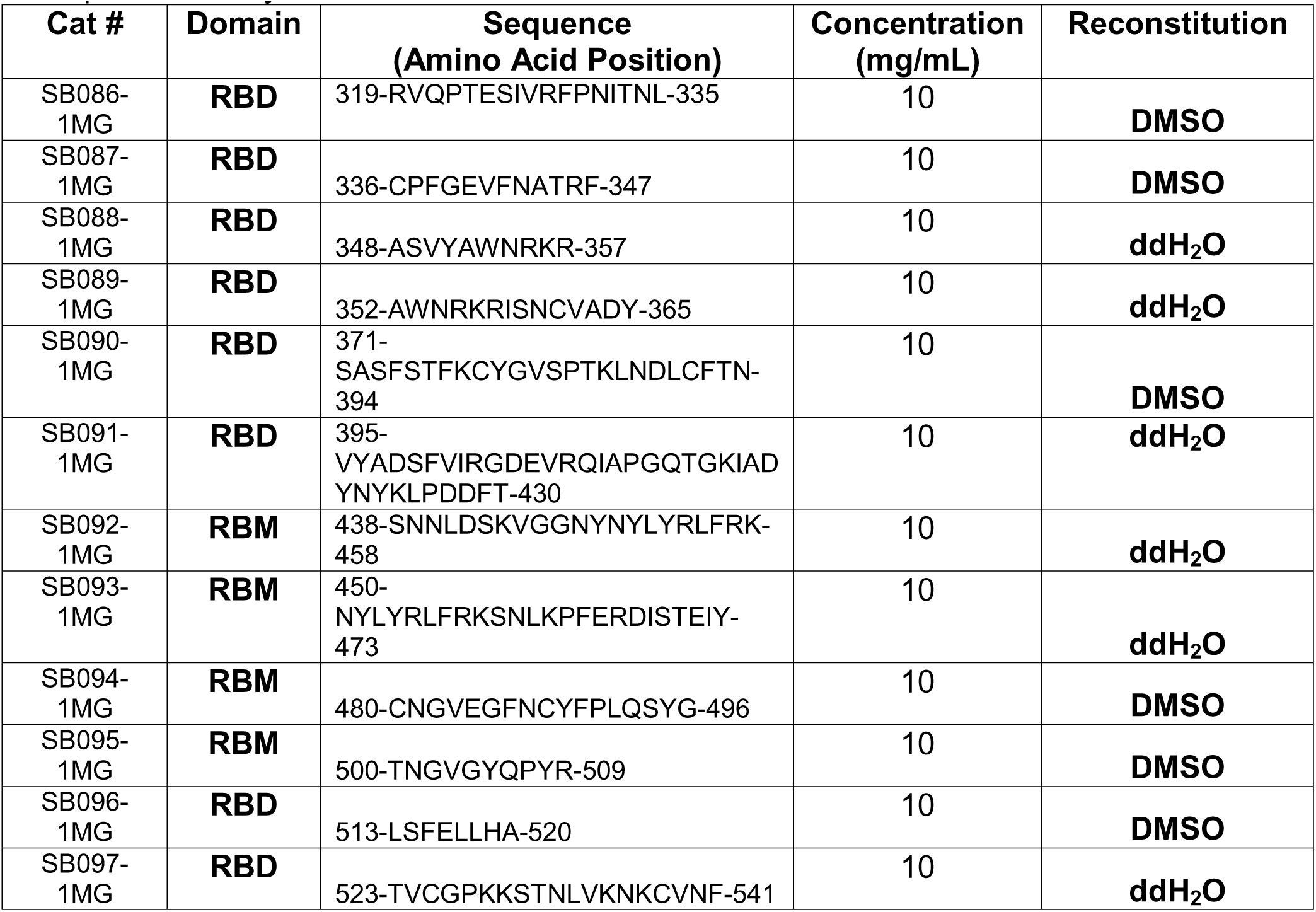
Synthesized SARS-CoV-2 Spike RBD Peptides for ACE2 Substrate Competition Assay.

### ACE2 Substrate Peptide Cleavage Competition Inhibition by ACE2 Inhibitor

The ACE2 activity assay was run as described above, with the addition of 100 µM MLN-4760 (Bio-Techne, Minneapolis, MN, Cat. #3345), a small-molecule ACE2 active-site inhibitor^55, 58^. This concentration is recommended in commercial ACE2 activity kits (Abcam, Waltham, MA. Cat. #ab273297).

### Enzyme Kinetics Assay

The ACE2 activity assay was set up as described, but with substrate dilutions ranging from 12.5 µM to 200 µM. An MCA standard curve (AAT Bioquest, Pleasanton CA. Cat. #557) was set up with 500, 250, 200, 150, 100, 50, 25, and 0 pmol MCA/well in 100 µL assay buffer. This curve enabled the calculation of pmol of MCA-APK(Dnp) substrate cleaved per minute.

### RBD-specific Binding Assays by ImmunoCAP

Following our previous studies^41, 42^, S-RBD binding was measured by using 50 µL of hybridoma supernatant with a quantitative ImmunoCAP system applying Phadia 250 (Thermo Fisher Scientific, Waltham MA. Thermo-Fisher/Phadia). Briefly, biotinylated S-RBD (RayBiotech, Peachtree Corners, GA) was conjugated to streptavidin-coated ImmunoCAPs (Thermo Fisher Scientific, Waltham MA. Thermo-Fisher/Phadia). Unconjugated streptavidin ImmunoCAP was run in parallel with its signal as a background to be subtracted from the singles of other samples.

### Immunoglobulin Depletion in Hybridoma Supernatants

Human immunoglobulin depletion was performed following pre-established methods ^41, 42^. Hybridoma culture supernatants were centrifuged at 12,000 × g at 4°C for 15 min, and supernatants were passed through a 0.45 µm syringe filter (FisherScientific, Cat. #97204). 500 µL of the filtered samples were added to ultrafiltration columns with a 100 kDa cutoff ultrafiltration membrane (Pierce Cat. # 88503) and centrifuged at 12,000 × g at 4°C until >400 µL of filtrate was produced. Then filtrated samples were incubated with 30 µL pre-washed Protein A/G Magnetic Beads (Life Technologies, Cat. # 88803, 10 mg/mL) at room temperature for 1 hour, followed by magnetic removal of the beads, followed by a second round of Protein A/G Magnetic Bead depletion with another freshly prepared aliquot of 30 µL magnetic beads. ACE2 substrate cleavage activity before and after immunoglobulin depletion was measured as described above. Immunoglobulin depletion efficiency was determined by human IgG ELISA, as described^41, 42^.

### Human IgG ELISA

The total amount of human IgG in hybridoma supernatant was measured by quantitative human IgG ELISA following the kit manufacturer’s instructions (Abcam, Waltham, MA. Cat. #ab100547) as described^41, 42^.

### Data Analysis

ACE2 substrate cleavage was monitored by measuring relative fluorescence units (RFU) for 16 hours with 5-minute intervals. The RFU values were corrected for background fluorescence by subtracting the minimum RFU value from the remaining RFU values for each well in the assay plate. Linear regression analysis was done on the first 5-120 minutes to determine the increase in RFU over time. This slope (RFU/min) was used as an indicator of ACE2-like activity. A slope of 0.5 RFU/min (the slope of 0.08 ng rhACE2) was determined to be the baseline detection level of ACE2 activity, which is two-fold greater than the lower limit of detection, the activity of 0.04 ng rhACE2^41, 42^. Monoclonal antibodies with a slope greater than 0.5 RFU/min were identified as positive for ACE2-like abzyme activity. For the MCA standard curve, the average RFU over 2 hours was plotted against pmol MCA/well. Linear regression was done to determine the number of RFU per pmol MCA. This value allowed for the calculation of pmols substrate cleaved per minute from the slope data. The supernatant IgG concentration was used to determine the *Kcat* of mAb catalytic activity. All data analysis and visualization were performed with R (version 2022-06-23) using R Studio user interface and packages including drc, ggh4x, ggplot2, patchwork, tidyverse, readr, GGally, corrplot and stringr.

## Results

We screened 129 purified human anti-RBD mAbs (Table 1) ^46, 47, 49, 50, 52, 59^ with one purified broadly neutralizing mAb against SARS-CoV-2 fusion peptide^51^ as an irrelevant control (Table 1). We also screened 10 human mAb hybridoma culture supernatants purchased from Precision Antibody, including 3 anti-RBD mAbs that blocked ACE2 binding to RBD, 4 anti-RBD mAbs that enhanced ACE2 binding to RBD, and 3 negative control irrelevant mAbs (Table 1). 5mM EDTA was added to the hybridoma supernatants to inhibit any native ACE2 protein in the media, following previous studies that inhibited ACE2 in 10% FBS media^56^. A concentration of 5 mM EDTA was confirmed to give complete inhibition of ACE2 in the hybridoma culture media serving as our negative control (FIG 1).

**FIG 1.**
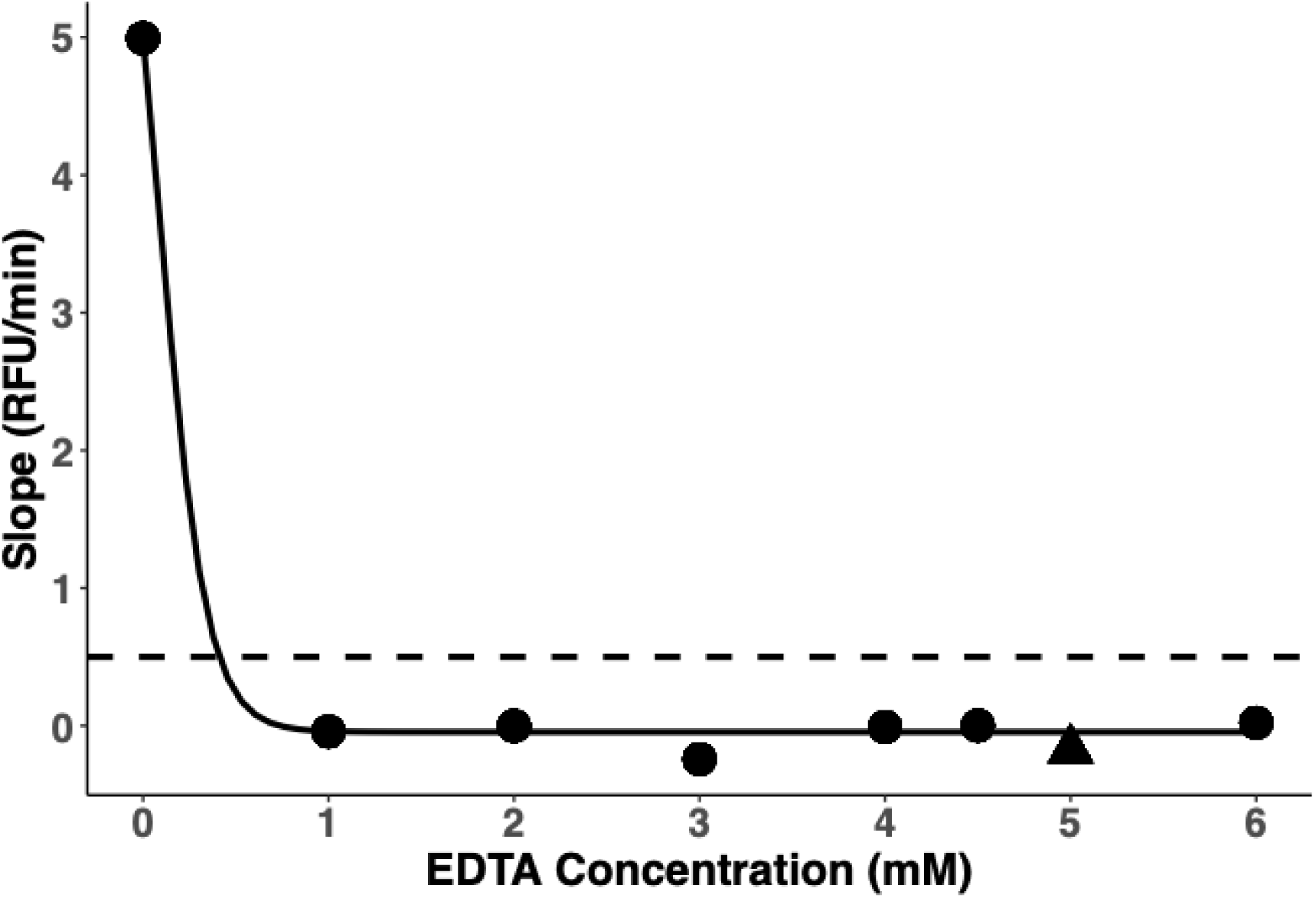
ACE2 Background Activity Inhibition by EDTA in Supernatant Control Media. ACE2 background activity in hybridoma supernatant control media (empty culture media without hybridoma) was detected by the ACE2 assay, indicated by the slope calculated from the kinetic curve. ACE2 background activity was effectively inhibited by adding EDTA to the media. The dashed line shows the cutoff, which was the limitation of the ACE2 assay and was determined using a purified recombinant human ACE2 positive control to call a sample positive for ACE2 activity. 5mM EDTA (**Triangle**) were used for following hybridoma culture supernatants screening as reported^56^.

We found 4 anti-RBD mAbs (AGRD013-4G9, AGRD016-2B3, AGRD016-4H2, AGRD021-9A10), that had ACE2-like abzyme activity (FIG 2. A). Two of these (AGRD013-4G9 and AGRD016-4H2) had previously been determined to block RBD binding to ACE2, while the other two (AGRD016-2B3 and AGRD021-9A10) had been determined to enhance binding of RBD to ACE2. An additional 9 mAbs had catalytic activity higher than the lower limit of detection, but lower than our predetermined threshold for significant ACE2 activity and thus were classified as negative for ACE2-like activity (FIG 2. B). All negative controls had slopes below the lower limit (FIG 2. B).

**FIG 2.**
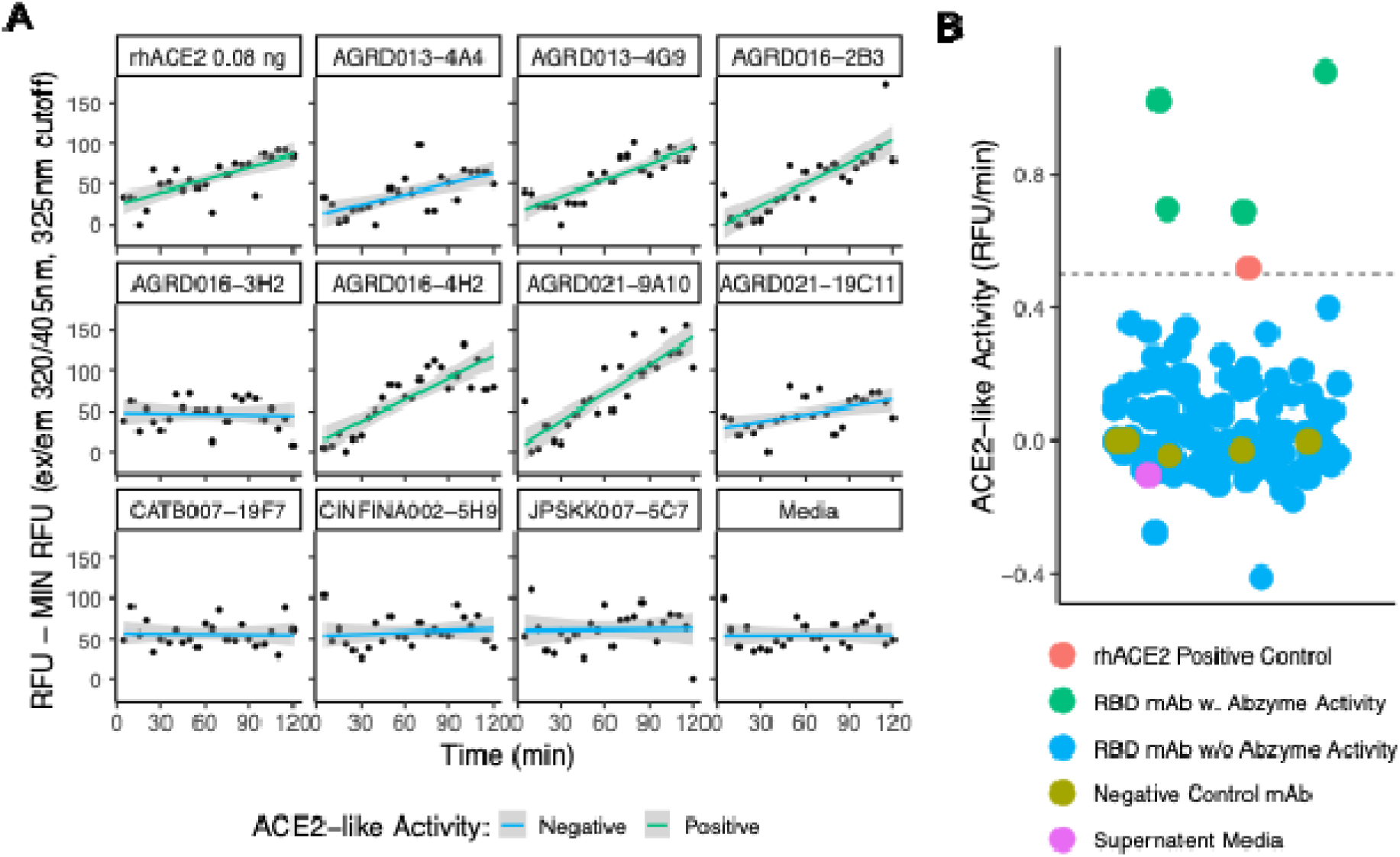
Initial Screening of mAbs for ACE2-like Activity. ACE2 substrate cleavage activity of mAbs. Catalytic activity was determined by calculating the slope of RFU from the initial 2 hours of the kinetics curve. 0.08 ng of purified recombinant human ACE2 served as a positive control and standard to define ACE2-positive activity. mAbs with ACE2-like activity greater than or equal to 0.08ng rhACE2 were identified as having ACE2-like catalytic activity. (A) Representative ACE2 activity screening kinetic curves from the Precision Antibody mAbs and ACE2 activity determination by slope. 0.08 ng rhACE2 served as a positive and standard control, and medium (empty hybridoma culture supernatant) served as a negative control. (B) Summary of average ACE2-like activity of all mAbs screened in this study. Results were from at least 2 rounds of screening. ACE2 substrate cleavage activity was determined by the slope of the RFU from the initial 2 hours of the kinetic curve. The dashed line shows the slope calculated from 0.08 ng of purified recombinant human ACE2, the initial 2-hour kinetic curve.

Antibody concentrations in hybridoma culture supernatants were measured by a total human IgG ELISA^41, 42^. The antibody levels ranged from 16 to 28 µg/ml. Anti-RBD binding activity in those hybridoma culture supernatants was assessed using an ImmunoCAP assay^41, 42^, with RBD antibody concentrations ranging from 8 to 13 µg/mL. Significant correlation (R = 0.92, *p* = 0.0035, Pearson correlation coefficient) was observed between total human IgG concentration and anti-RBD IgG (FIG 3), indicating that ACE2-like activity correlates with anti-RBD antibody, consistent with our previous findings^41, 42^.

**FIG 3.**
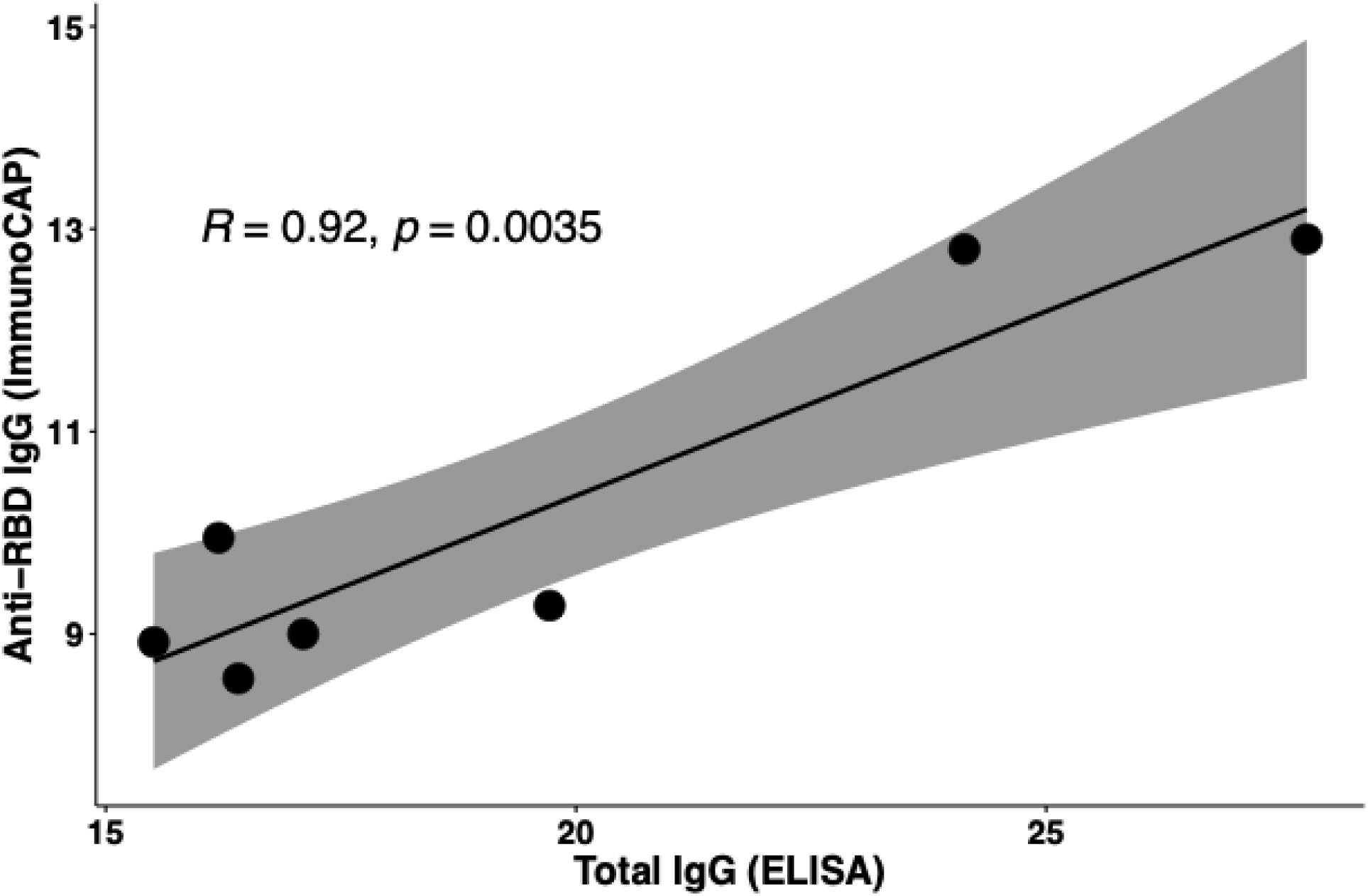
Correlation of total IgG and anti-RBD antibody. Total IgG concentrations in mAb hybridoma culture supernatants were measured by human IgG ELISA. Anti-RBD IgG concentrations of mAb hybridoma culture supernatants were determined by ImmunoCAP antibody-binding assay. (Pearson correlation test).

Studies of the enzymatic activities of those mAbs, ACE2-like activity positive samples, and the rhACE2 positive control, was conducted using Michaelis–Menten and Lineweaver-Burk analyses^60^. IgG concentrations were determined by human IgG ELISA (FIG 3, Table 3). The ACE2-like abzymes have lower *K_m_s*, and ∼4 log_10_ lower catalytic efficiency (Table 3, FIG 4). The low catalytic efficiency (*K_cat_/K_m_*) is consistent with the data obtained for other abzymes^15, 16^. Our calculated *K_cat_/K_m_* for rhACE2 at 37°C is 0.8 M^-1^s^-1^ higher than the reported value for rhACE2 at room temperature^61^.

**Table 3:** Enzyme Kinetic Studies of ACE2-like Abzymes and Purified Recombinant Human ACE2 Molecule (From FIG 4)

| Sample | Assay Conc | $K_m$ ($\mu M$) | $V_{max}$ (pmol/min) | $V_{max}$ (mol/s) | $K_{cat}$ ($s^{-1}$) | $K_{cat}/K_m$ ($M^{-1}s^{-1}$) |
| --- | --- | --- | --- | --- | --- | --- |
| rhACE2<sup>a</sup> | 25 ng/mL | 43.7 | 66.9 | 1.12E-12 | 3.79E+01 | 8.68E+05 |
| AGRD013-4G9 | 5.8 $\mu g/mL$ | 26.9 | 0.192 | 3.20E-15 | 8.27E-04 | 3.07E+01 |
| AGRD016-2B3 | 5.6 $\mu g/mL$ | 34.3 | 0.147 | 2.45E-15 | 6.57E-04 | 1.91E+01 |
| AGRD016-4H2 | 7.9 $\mu g/mL$ | 38.9 | 0.295 | 4.92E-15 | 9.33E-04 | 2.40E+01 |
| AGRD021-9A10 | 5.3 $\mu g/mL$ | 33.9 | 0.253 | 4.22E-15 | 1.19E-03 | 3.52E+01 |

a: Purified recombinant human ACE2 molecule as positive control.

**FIG 4.**
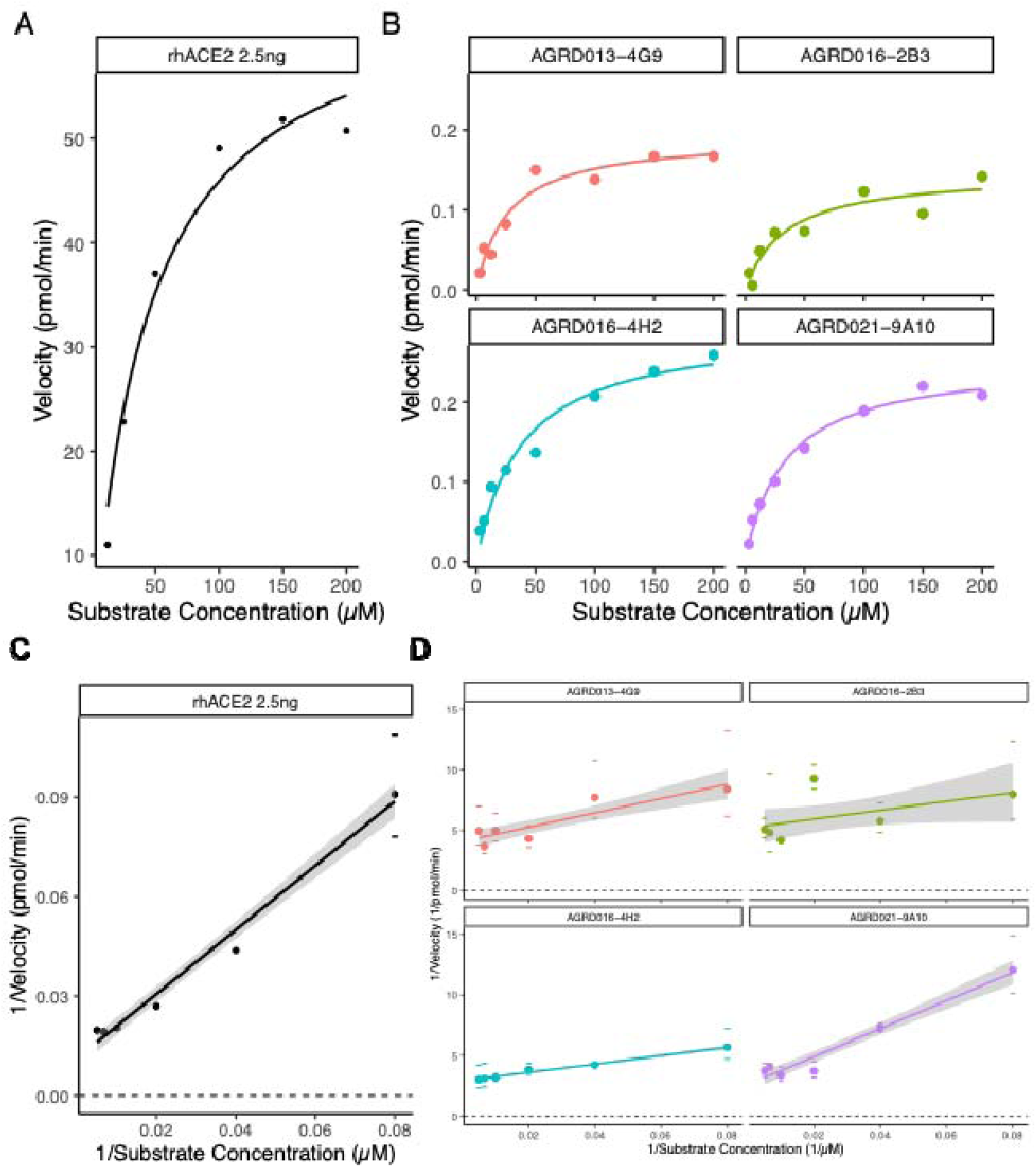
Michaelis-Menten and Lineweaver-Burk Analysis of mAb Catalytic Activity. Michaelis-Menten plots for (A) Positive control (2.5ng rhACE2) and (B) mAbs that had fluorogenic ACE2 surrogate peptide substrate cleavage activities. Lineweaver-Burk plot for (C) Positive control (2.5ng rhACE2) and (D) mAbs that had ACE2 substrate cleavage activities. Abzyme IgG amount for catalytic kinetics was from human IgG ELISA.

Substrate preference of abzyme catalytic activity was determined by comparing the cleavage of ACE2 substrate (MCA-APK(Dnp)) and control substrate (MCA-Pro-Leu-OH), as in prior studies^41, 42, 62^. In contrast to robust signal generation with the ACE2 substrate, reactions containing the control substrate produced negative slopes (FIG 5), consistent with the absence of measurable cleavage. These findings indicate that the antibodies with identified catalytic activity preferentially cleaved the ACE2 substrate rather than nonspecifically cleaving the control substrate.

**FIG 5.**
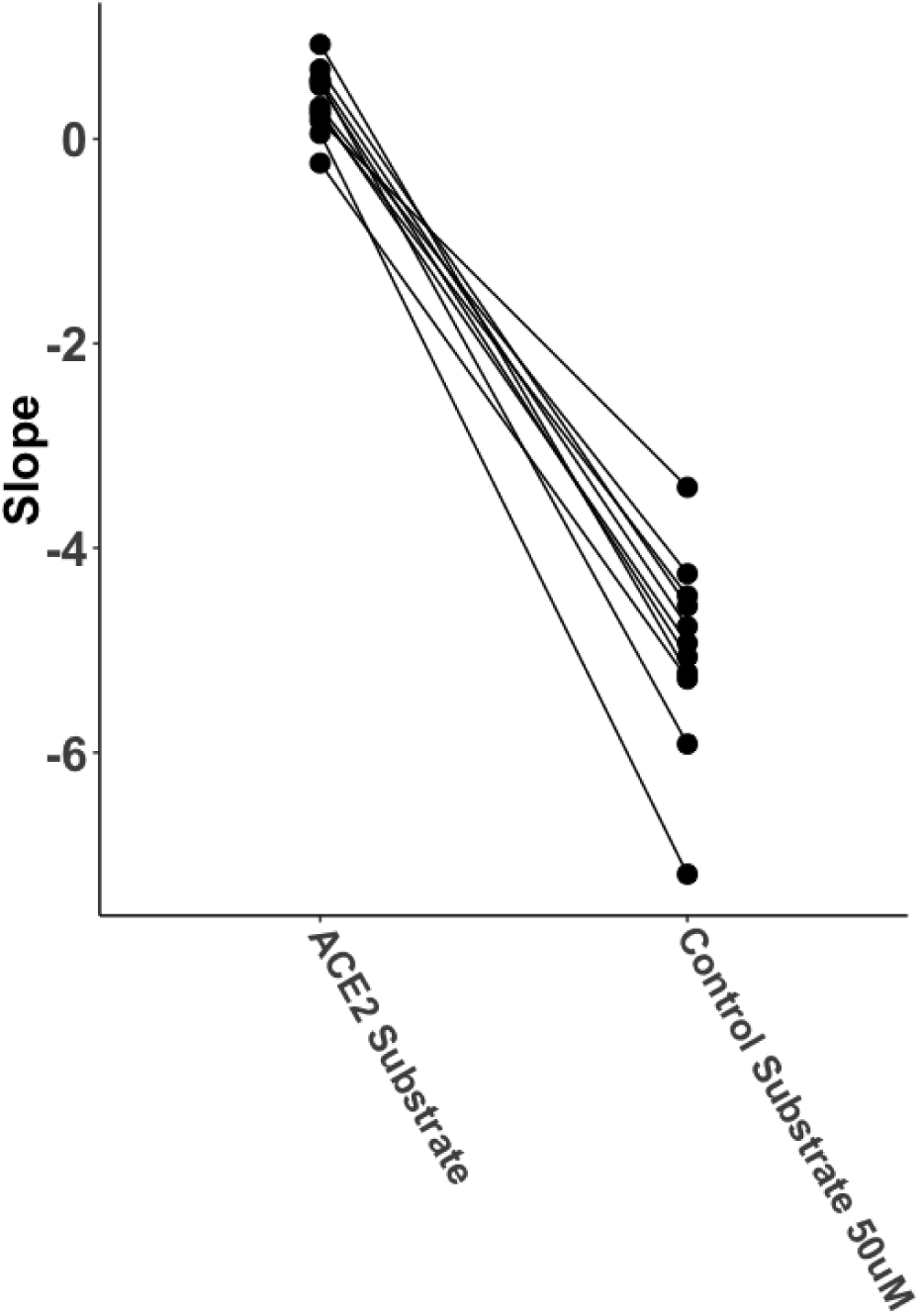
Comparative Substrate Preference of Abzymes. The substrate cleavage activity of the mAbs was characterized by comparing using a fluorogenic ACE2 peptide substrate, MCAIZlAPK(Dnp), and a control non-cleavable substrate. The slope of the RFU from the initial 2 hours of the kinetic curve was calculated to determine ACE2-like activity. No catalytic activity was observed with the control substrate indicated by negative slope.

To further confirm the catalytic activity of the monoclonal antibodies, we conducted ACE2 substrate cleavage peptide inhibition assays using 12 tiled overlapping peptides that cover the SARS-CoV-2 RBD and RBM amino acid sequences (Table 2). The RBD peptide pools significantly inhibited the ACE2-like activity of two mAbs, AGRD013-4G9 and AGRD016-4H2, which block RBD binding to ACE2, but had overall limited inhibition effect on two other mAbs, AGRD016-2B3 and AGRD021-9A10, which enhance RBD binding to ACE2 (FIG 6). The RBD peptide inhibition results show that the mAb catalytic activity could be inhibited by peptide competition, and that there is a difference in the ability of the peptides to inhibit activity between mAbs which block or enhance receptor binding^41, 42^.

**FIG 6.**
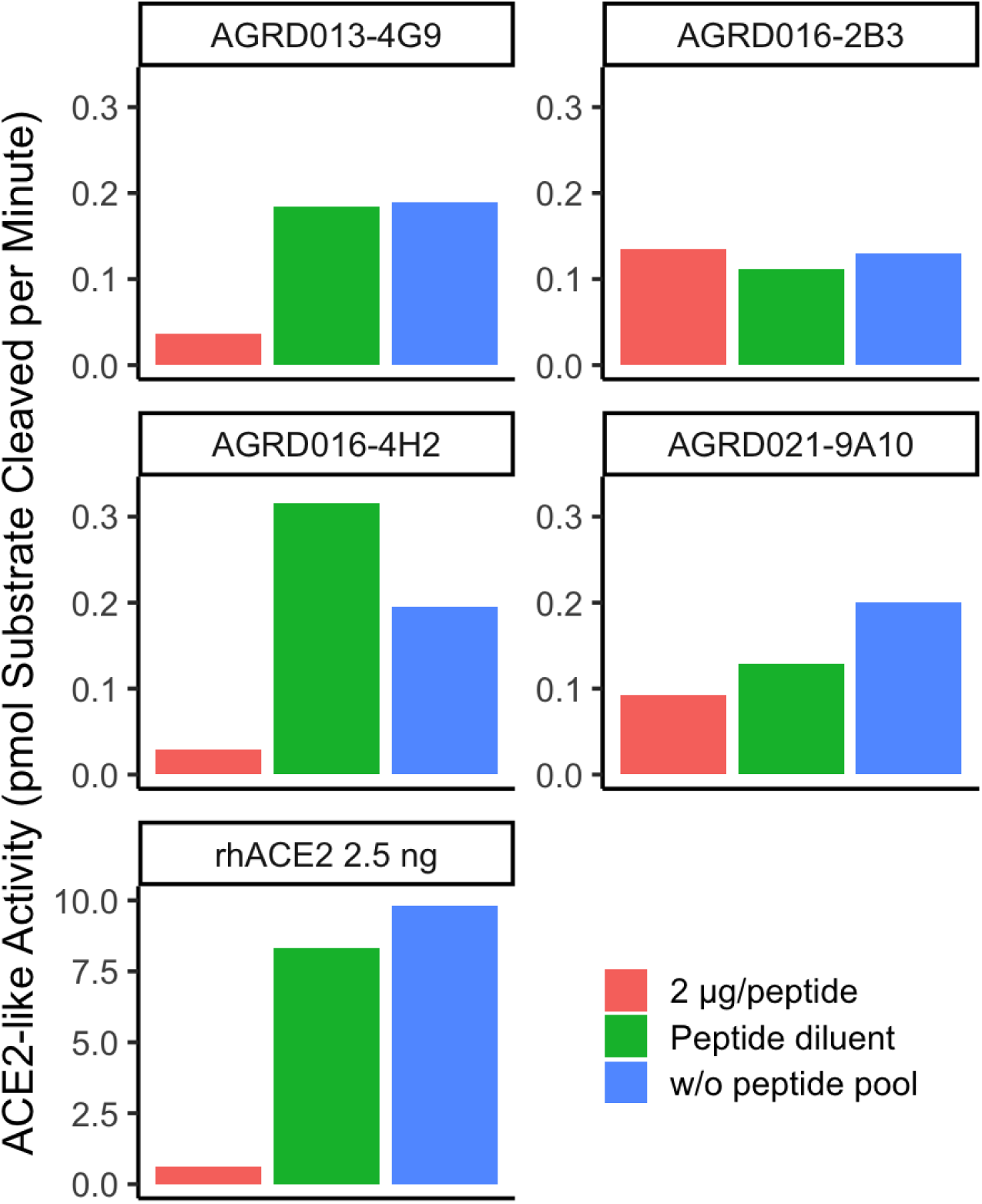
SARS-CoV-2 RBD Peptide Competitive Inhibition of mAb ACE2 Substrate Cleavage Activity. mAbs ACE2-like activity inhibition by RBD peptide competition was performed by adding RBD peptide pools to ACE2 assay reactions containing mAbs defined as ACE2-like activity-positive. Peptide pool diluent, with an equal volume of peptide pool added to the assay reactions, served as a comparison. 2.5 ng purified recombinant human ACE2 was used as a positive control. ACE2-like activity was determined by calculation of the slope of the RFU from the initial 2 hours of the kinetic curve.

To confirm the ACE2 substrate cleavage activity was immunoglobulin related, we completely depleted the immunoglobulin from mAb hybridoma culture supernatants (depletion efficiency >99.99%, Table 4) by previously established IgG depletion assay ^41, 42^. Elimination of ACE2 substrate cleavage activity was observed with immunoglobulin depletion (FIG 7) indicating the ACE2 substrate cleavage activity is associated with immunoglobulin.

**Table 4:**
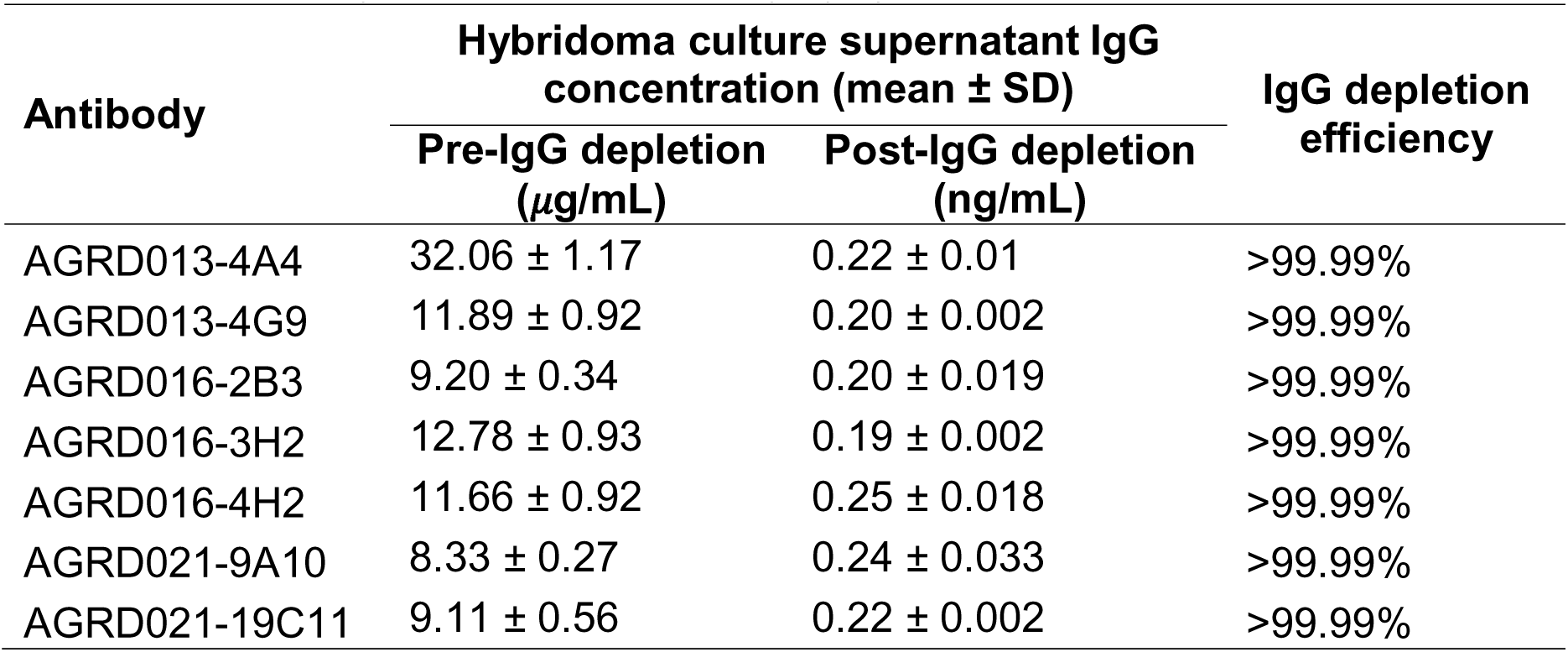
Human IgG Depletion Efficiency by IgG ELISA.

**FIG 7.**
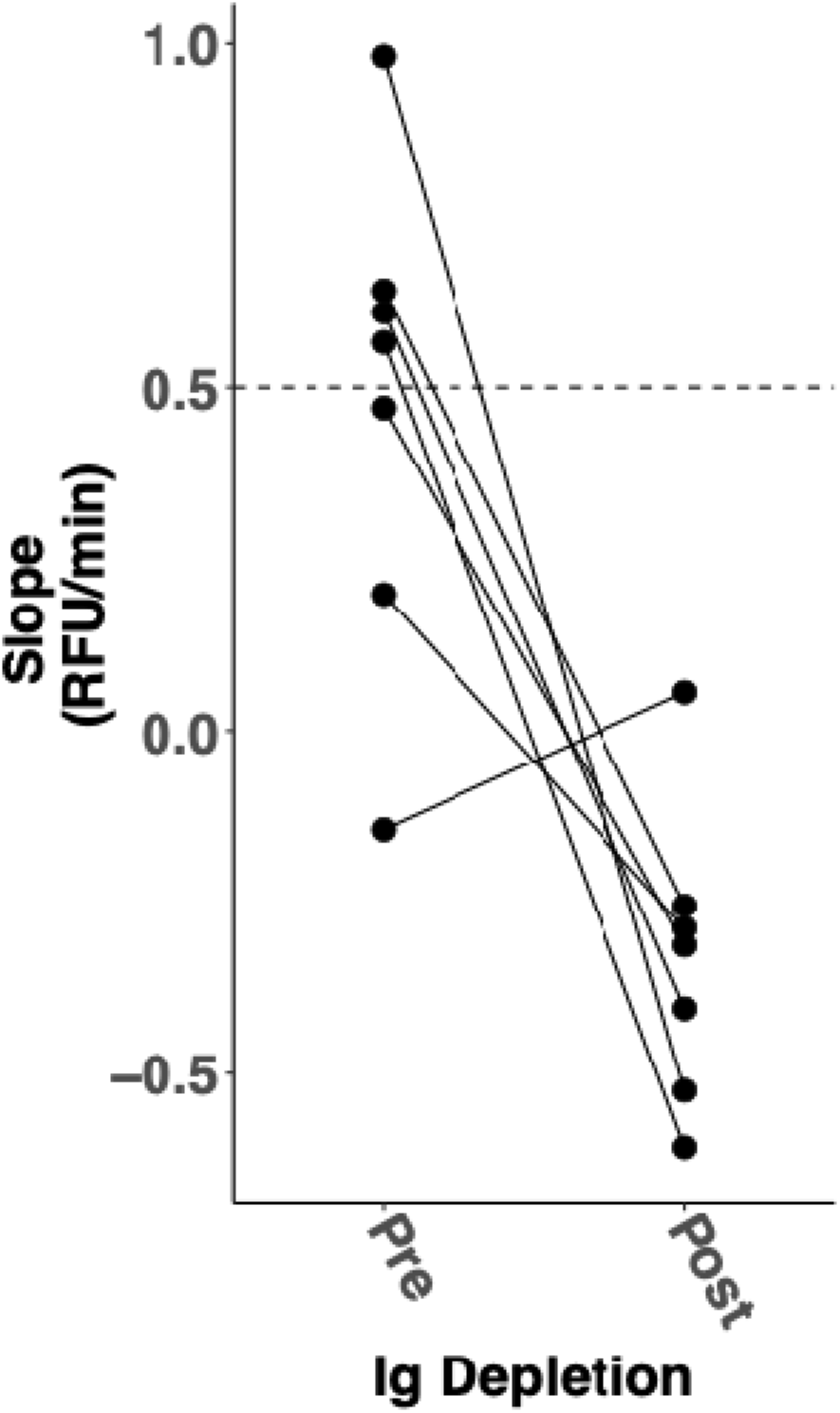
ACE2 Substrate Cleavage Activity of mAb Hybridoma Culture Supernatants before and after Immunoglobulin Depletion. ACE2 substrate cleavage activities of mAb hybridoma culture supernatants before (Pre) and after (Post) immunoglobulin depletion were parallel compared. The slope of the RFU from the initial 2 hours of the kinetic curve was calculated to determine ACE2-like activity. The dashed line shows the cutoff, which was the limitation of the ACE2 assay and was determined using a purified recombinant human ACE2 positive control to call a sample positive for ACE2 activity. IgG depletion efficiency was determined by human IgG ELISA (Table 4). No catalytic activity was observed from IgG complete depleted samples indicated by negative slope of kinetic curve.

We also tested the effect of a small molecule ACE2 inhibitor, MLN-4760, a well-documented inhibitor of ACE2 catalytic activity^55, 58^. MLN-4760 completely inhibits rhACE2, consistent with previous findings^55, 58^, but did not inhibit ACE2-like abzyme activity (FIG 8). As we previously showed, the divalent cation chelator, EDTA did not affect the mAbs’ substrate cleavage activity, but inhibited native ACE2 catalytic activity, since native ACE2 is a Zn^2+^ metalloprotease^41, 42^. These findings suggest that the mAbs with ACE2-like activity employ different catalytic mechanisms than native ACE2.

**FIG 8.**
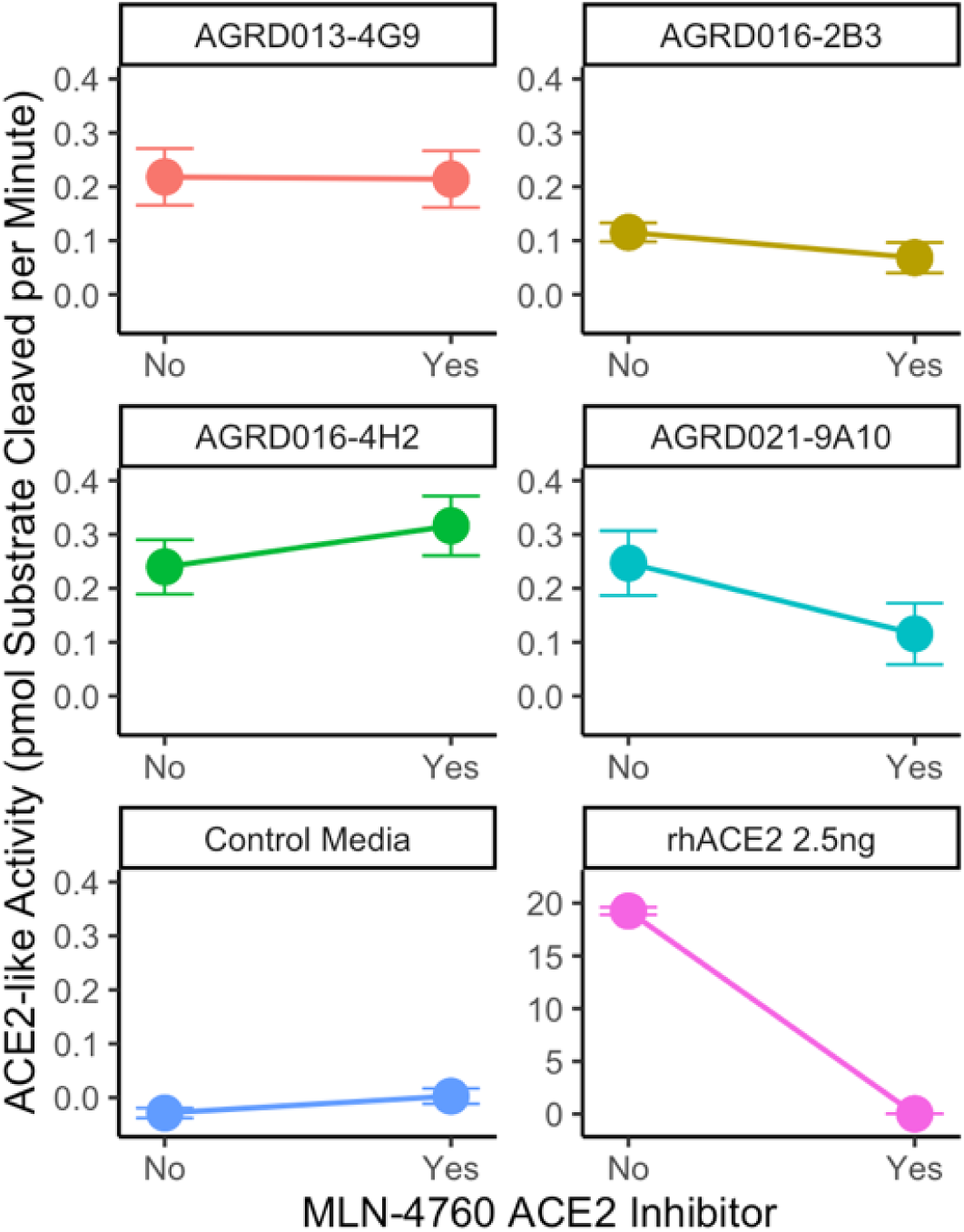
Effect of ACE2 Inhibitor, MLN-4760, on ACE2 Substrate Cleavage Activity of mAbs and rhACE2. 100 M ACE2 inhibitor, MLN-4760, was added to ACE2 assay reactions for comparison of ACE2 activity with ACE2 assay reactions that had no inhibitor. 2.5 ng purified recombinant human ACE2 was the positive control, and control media (empty hybridoma culture supernatant) was the negative control. ACE2-like activity was determined by calculation of the slope of the RFU from the initial 2 hours of the kinetic curve.

## Discussion

The COVID-19 pandemic has claimed around 7 million lives, according to the World Health Organization, as of January 2026^63^, with about 65 million patients experiencing lingering symptoms long after their initial infection^3^. Long COVID, characterized by persistent symptomatology following an initial COVID-19 infection, includes symptoms consistent with dysregulated proteolytic regulatory cascades^29–31^ such as the angiotensin-renin system ^38^.

Our previous research showed that some patients suffering from acute SARS-CoV-2 infection or convalescing from COVID-19 produced antibodies with ACE2-like catalytic activity ^41, 42^. In this study, to further explore our hypothesis that some anti-spike antibodies exhibit ACE2-like catalytic activity, we screened 129 anti-SARS-CoV-2 mAbs, 7 anti-spike mAb hybridoma supernatants; and 3 irrelevant mAb hybridoma supernatants and an anti-SARS-CoV-2 FP mAb as controls (Table 1). Although most anti-RBD mAbs exhibited no ACE2-like activity, we identified 4 anti-RBD mAb hybridoma supernatants that exhibited measurable catalytic activity against a synthetic fluorogenic peptide substrate commonly used to assess metalloprotease function. This finding is consistent with our hypothesis that only a small fraction of elicited RBD antibodies exhibit a widely used surrogate ACE2 substrate cleavage activity after infection underscoring the rarity of this phenomenon ^24, 25^.

The catalytic activity of metalloprotease ACE2 is inhibited by 5 mM EDTA in the assay buffer. The ACE2-like catalytic activity of the mAbs was not inhibited by 5 mM EDTA (FIG 1), just as we observed directly in samples from research volunteers with both acute COVID-19 and convalescing from COVID-19^41, 42^. ACE2 substrate cleavage was further confirmed by conducting assays using either a native substrate or a negative control ACE2 substrate widely used in established ACE2 activity detection assays (FIG 5)^41, 42, 62^, and a SARS-CoV-2 spike peptide pool in a competition/inhibition assay (FIG 6), which has been used to demonstrate immunoglobulin-associated ACE2-like activity in clinical samples^41, 42^. The peptide pool inhibited the substrate cleavage of two mAbs (AGRD013-4G9 and AGRD016-4H2), which blocked ACE2 binding to the virus spike protein, suggesting that the catalytic sites of the mAbs have sufficient resemblance to the ACE2 to be inhibited by the overlapping spike-derived peptide pool. However, two mAbs (AGRD016-2B3 and AGRD021-9A10) enhance spike protein binding to ACE2, indicating the mAbs have distinct structural and functional features. ACE2 substate cleavage activity was removed after IgG complete depletion (Table 4 and FIG 7) further confirmed the association of catalytic activity with immunoglobulin.

The four mAbs with ACE2-like activity exhibit catalytic efficiency (Table 3 and FIG 4) 4 log_10_ lower than human ACE2 molecule, a known characteristic of other abzymes^16^, so the characteristics of the isolated mAbs with ACE2-like catalytic activity fit well within those described for other abzymes. The inability of the native ACE2-specific small-molecule inhibitor, MLN-4760, to inhibit the ACE2-like activity of abzymes (FIG. 8) suggests that the active site structures of the mAbs differ from those of native ACE2^12, 13, 17^. These experiments contribute to further characterizing the magnitude and biochemical properties of antibodylilassociated catalytic activity.

Our study had several strengths. We screened many mAbs from four sources for ACE2-like catalytic activity. Although a large fraction of the tested mAbs did not have ACE2-like catalytic activity, we found that four isolated anti-SARS-CoV-2 spike mAbs have ACE2-like catalytic activity. Our previous studies describing immunoglobulin-associated ACE2-like activity in clinical samples could have had other, unappreciated substances, other than Abs, responsible for the ACE2-like catalytic activity. The identification of ACE2-like catalytic activity in mAbs uncontaminated by the many substances present in clinical plasma samples strongly supports the existence of Abs with ACE2-like catalytic activity.

It is important to note that the Spike RBD binds to a region on ACE2 that does not physically fully overlap with its catalytic cleft^64^. This suggests that the ACE2-like activity we observed is likely not a result of a complete mimicry of the ACE2 active site. It is more likely that some Spike components may structurally resemble the transition state of ACE2-mediated peptide hydrolysis. Consequently, antibodies templated by these Spike motifs, along with the regions that bind ACE2, may possess adventitious catalytic pockets—a phenomenon previously observed in other autoimmune pathologies such as systemic lupus erythematosus (SLE) and Hashimoto’s thyroiditis^18, 65^.

However, our study had certain limitations. By screening pre-established mAb collections from our collaborators^46, 49–52, 59, 66^, the mAbs were produced without clear linkage to patients with symptoms associated with COVID-19. In particular, the mAbs were not produced from LC patients with some of the most problematic symptoms. This prevents correlation of abzymes characteristics with clinical symptoms. In addition, the motivation behind the production of most of the mAbs was to obtain mAbs capable of inhibiting SARS-CoV-2 infection of cells. Selecting for a mAb that is highly effective at blocking the virus from infecting a cell may bias the mAb production process away from selecting mAbs with catalytic activity, since those mAbs may not be particularly effective at blocking cell infection. In patients, highly neutralizing Abs may not dominate the total pool of Abs made in response to SARS-CoV-2 infection^67, 68^.

All four mAbs with ACE2-like activity were identified in hybridoma culture supernatants from a single source; the 129 purified mAbs from the other three sources were showed no ACE2-like catalytic activity. Several hypotheses, which are not mutually exclusive, could account for this pattern. The most straightforward is that differences in screening and selection methodology enriched the Precision Antibody panel for clones predisposed to abzyme activity. Those mAbs were specifically selected for their ability to modulate ACE2 binding to the spike RBD — a criterion that favors isolation of antibodies whose combining sites are structurally complementary to the ACE2-RBD interface, which is precisely the structural prerequisite for ACE2-like catalytic activity.

The mAbs from the other three sources were selected under different criteria, primarily neutralization potency or epitope coverage, which optimize for different features, which may be less likely to include catalytic sites. A second, independent hypothesis is that mAb purification procedures disrupted catalytic activity in the other panels. Low-pH Protein A/G elution steps can alter antibody conformation, and a catalytic pocket, which depends on precise active-site geometry, may be more conformationally labile than an antigen-binding site, which is governed primarily by CDR contacts that are generally robust in purification. Finally, the differences in panel size and format across sources mean that statistical sampling cannot be excluded as a partial contributor — the probability of detecting rare abzyme-positive clones is lower in small panels, and is further reduced if purification suppresses activity. Importantly, none of these considerations undermine the validity of the positive supernatant results, which satisfy the same criteria used in our prior clinical studies: the activity co-depletes with immunoglobulin at greater than 99.99% efficiency and is inhibited by competition with a tiled overlapping RBD peptide pool, establishing that it is antibody-associated and antigen-specific. The ACE2-like catalytic activity of the hybridoma supernatants did not require divalent cations (was not inhibited by EDTA), so the activity is distinct from native ACE2. The low catalytic efficiency observed here is consistent with the broader abzyme literature, in which antibody associated catalysis typically displays slow turnover, modest substrate affinity, and considerable heterogeneity^22, 69–71^. Previous well-documented abzymes from autoimmune diseases like Systemic Lupus Erythematosus (SLE) and Multiple Sclerosis exhibited significantly lower catalytic efficiency than that of dedicated nucleases or proteases, yet the abzymes remain fundamentally pathogenic and leading to chronic tissue damage due to their ability to work under physiological conditions^65, 72^. Future studies should prioritize production and testing of mAbs from LC patients not selected for neutralization activity, evaluated in both supernatant and purified formats under varying purification conditions, to systematically disentangle these hypotheses.

Native enzymes are strictly regulated by localization and inhibitors, in contrast, IgG-based abzymes circulate with a long biological half-life and lack the features that help control the physiological regulatory cascade pathways like the renin-angiotensin system^20, 22, 23^. The cumulative effect over months of disease progression as occurs in LC or autoimmune diseases could be substantial If these abzymes similarly cleave native substrate under physiological condition, even at a fraction of the rate of native enzymes^73–76^. In a patient with high titers of these catalytic IgGs, the constant, unregulated processing of signaling peptides leading to a significant integrated deficit in homeostatic peptide levels over months could contribute to the vascular and inflammatory dysregulation frequently observed in Long COVID^77^ ^34^.

The data presented here established that catalytic activity can be detected in rare anti-Spike antibodies under defined *in vitro* conditions using a highly sensitive model substrate. In future studies, to detect these low catalytic activities it may be important to ensure that the antibodies are maintained in an physiological intact state during isolation, since it is conceivable that binding or infection inhibition may be less fragile properties that catalytic activity^78–81^, which could be a conceivable reason why catalytic activity was not detected in the purified mAbs. Nevertheless, the identification of any mAbs with ACE2-like catalytic activity supports the hypothesis underlying this study.

Another limitation of this work is that catalytic activity was assessed using a short synthetic fluorogenic peptide, Mca-APK(Dnp), rather than a native biological substrate. While this synthetic substrate is cleaved by ACE2 and is useful for detecting low level metalloprotease like activity, it differs substantially from endogenous ACE2 substrates such as angiotensin II in size, sequence context, and structural constraints ^82^. This substrate was specifically engineered to mimic the C-terminal sequence of natural ACE2 substrates and has been validated as a highly sensitive and selective surrogate for ACE2 carboxypeptidase activity^44, 45^. Mca-APK(Dnp) hydrolysis has been shown to follow the same zinc- and chloride-dependent mechanisms as the natural substrate, Angiotensin II^83, 84^. Its use in this study reflects its established utility as a sensitive surrogate for real-time and continuous kinetic monitoring ACE2lilclass metalloprotease activity, which is essential for the precise determination of *Vmax* and *Kcat* in low-activity antibody preparations where endpoint-based HPLC methods may lack sufficient sensitivity^84–90^. We noted that these analyses were designed to assess substrate preference rather than absolute specificity and did not exclude the possibility that additional substrates may also be cleaved at lower efficiency. A comprehensive definition of substrate specificity would require systematic testing across diverse peptide libraries and protein substrates in future studies, which was beyond the scope of this study.

An essential next step involves elucidating the mechanistic properties of the abzymes’ catalytic activity. Structural analyses aimed at delineating these activities could provide critical insight into the mechanisms of ACE2-like catalytic activity. A clear understanding of the catalytic mechanisms of the mAbs with ACE2-like catalytic activity could enable the development of inhibitors against that activity, which would represent a potential novel avenue for LC therapeutics development. Alternatively, the source of pathologic antibodies, the cells producing the pathologic abzymes, could be targeted.

Overall, our study further confirms the existence of anti-spike antibodies with ACE2-like activity that can plausibly contribute to the pathogenesis of COVID-19 and LC. While much work remains to be done, including the production and study of new anti-spike mAbs not selected specifically for SARS-CoV-2 neutralizing activity, structural studies of mAbs with catalytic activity, and the investigation of catalytic activities other than ACE2 in Abs made after exposure to SARS-CoV-2 that could potentially mediate the pathogenesis of COVID-19 and LC. Additional studies in understand COVD-19 an dLC pathogenesis would include work aimed at associating different clinical features of disease with abzymes that may have particular characteristics, structural and mechanistic studies of the abzymes themselves, and larger epidemiologic survey studies to better determine the rates of abzyme production in persons exposed to SARS-CoV-2.

There are also longer-term translational implications. Other viruses, in addition to SARS-CoV-2, use cell surface enzymes as their receptors ^91–96^. If pathologic abzymes induced by exposure to SARS-CoV-2 are responsible for some of the pathologic features of COVID-19 and LC, then abzymes induced by other viruses, including viruses causing subclinical infections, could be responsible for other disorders.

## Institutional Review Board Statement

The UVA IRB approved enrollment of all subjects and collection of specimens and related metadata (HSR #HSR13166).

## Informed Consent Statement

Informed consent was obtained from the subjects in the study.

## Data Availability Statement

Data is presented in the manuscript. Raw data is available upon request.

## Acknowledgments and Funding

The study was supported by University of Virginia internal funding, including the Pendleton Laboratory Fund for Pediatric Infectious Disease Research, the Manning Fund for COVID-19 Research at UVA, the Ivy Foundation, University of Virginia Summer Research Internship Program and grants from NIAID, NIH (R01 AI176515). Some data for this study were generated using instrumentation supported by the UVA Comprehensive Cancer Center (P30-CA044579). The authors acknowledge Research Computing at The University of Virginia for providing computational resources and technical support that have contributed to the results reported within this publication. URL: https://rc.virginia.edu. This research was supported in part by the Intramural Research Program of the National Institutes of Health (NIH). The contributions of the NIH author(s) were made as part of their official duties as NIH federal employees, are in compliance with agency policy requirements, and are considered Works of the United States Government.

However, the findings and conclusions presented in this paper are those of the author(s) and do not necessarily reflect the views of the NIH or the U.S. Department of Health and Human Services. AI agents (Claude, Copilot, Edison Scientific, Gemini) were use code-writing, debugging, and optimization; literature searches and finding source identification, manuscript editing, cross-checking, and proof-reading. We thank Peter Kwong, Columbia University, Aaron Diamond AIDS Research Center, for helpful advice and comments.

